# Tropism of Drosophila innubila Nudivirus in Cell Culture and in Tissues

**DOI:** 10.64898/2026.10.12.758614

**Authors:** Kent M. Mulkey, Margaret E. Schedl, Brian D. Ackley, Robert L. Unckless

**Author notes:** Address correspondence to Robert L. Unckless,.

## Abstract

Most viruses are successful because they can infect and exploit specific host tissues for reproduction and transmission. As a result, we often classify viruses by their ability to infect different tissues: respiratory, reproductive and gut, for example. Thus, knowing the tissue and cellular tropism of a virus may elucidate aspects of its biology including routes of transmission and virulence. We studied tropism of the *Drosophila innubila* Nudivirus (DiNV) in cells and infected hosts. DiNV is a nudivirus, a family of viruses related to the baculoviruses that are relatively large (150-200 nm), contain large double-stranded DNA genomes, and infect different host tissues. Cultured cells stained with an antibody against the DiNV nucleocapsid demonstrated punctate staining in the cytoplasm by confocal microscopy. Electron micrographs of infected cells, on the other hand demonstrated that virions are present in the nucleus. After purification, DiNV was often associated with multimembrane-like vesicles as described for Oryctes rhinoceros nudivirus. We examined tropism in hosts first by sagittal section and found most virus associated with the gut and, to a lesser extent, reproductive tissues. Upon dissection and electron microscopy, we found that DiNV in the gut is primarily in the visceral muscle of the gut and not the lumen as might be expected for a virus if transmitted fecal-orally.

## Introduction

Most viruses are successful because they can infect and usually exploit specific host tissues which enhances their transmission and ultimately fitness (1, 2). As such, viral tissue and cell tropism is well studied for many of the viruses most important for human and livestock health (3, 4). Respiratory viruses such as SARS-CoV-2, influenza, and Respiratory Syncytial Virus (RSV) primarily, though not exclusively, infect the airways (5-7). Hepatitis viruses primarily infect the liver, and enteroviruses (poliovirus, coxsackie virus) primarily infect the gut and associated tissue (8-12). Understanding the tissue tropism of these viruses contributes to our understanding of their life cycles, pathogenesis and treatment (13, 14). It is therefore somewhat surprising that we know very little about tissue tropism for many of the excellent model host/virus systems in arthropods. While the body plans of arthropods are simpler in many ways, they have structures that are homologous to those found in mammals so understanding tissue tropism should inherently inform the dynamics of the model system.

*Drosophila* have served as a model system to study everything from space flight to cancer to alcoholism (15-17). They have proven particularly powerful as a model for studying host/pathogen interaction and immunity because of the parallels between *Drosophila* and human innate immunity (18-20). While RNA viruses have been part of this effort for decades, DNA viruses that naturally infect *Drosophila* in the wild were notoriously absent until the 2010s (21-25). Still the most numerous DNA viruses in wild *Drosophila* are a handful of nudiviruses, a class of viruses related to the baculoviruses but largely lacking the characteristic occlusion body (which earned them their libertine name, “nude”). Nudiviruses often infect both larval and adult stages and their peak virulence differs among viruses. Oryctes rhinoceros Nudivirus (OrNV) is most virulent in larvae and has been used in pest control for palm beetles since the 1970s (26, 27). Interestingly, there are two different nudiviruses that infect the corn earworm, *Helicoverpa zea* (28). The first, *Helicoverpa zea* Nudivirus-1 (HzNV-1) is unable to infect larvae and has only been observed in cell culture (29). HzNV-2, on the other hand, infects larvae, leads to gonadal atrophy and can be transmitted vertically through the mother or horizontally *via* sexual contact (30). The two viral genomes are quite similar with 99% nucleotide identity though HzNV-2’s genome is about 3000 bases larger (228089 vs 231621 bp) (28). Across nudiviruses, horizontal transmission including fecal-oral seems to be the most common route of infection and, as a result, guts and fat bodies tend to be the most commonly infected tissues (28). So, there is precedence for nudiviruses infecting gut, immune (fat body) and reproductive tissue with varying consequences.

The best studied of the *Drosophila* nudiviruses are Kallithea virus, which was discovered in *D. melanogaster*, and Drosophila innubila Nudivirus (DiNV), which was discovered in *D. innubila* (31-34). Both are associated with increased mortality and reduced female fecundity and encode a suppressor of host NF-KappaB signaling that likely enhances virulence (31, 34). While it is unknown whether these viruses directly infect the ovaries, the reduced fecundity paired with ovarian atrophy upon infection with Kallithea virus (32) suggest that reproductive tissues may be a target of infection. There is also indirect evidence for gut infection in DiNV since it was observed to pass from individual to individual within a vial *and* was found in infected-fly feces (22). Finally, there is little evidence for vertical transmission of DiNV through females (22). We were recently able to culture the virus in cells and fluids recovered from that cell culture are able to infect and kill *D. innubila* 7-10 days after infection (35, 36).

We determined the tropism of DiNV within the cell and throughout the animal after virus injection. We focus on systemic infection through injection because we are better able to control dose and because, to date, we have been unable to establish infection through feeding with our cell-culture adapted virus. We find DiNV in multimembrane-like vesicles in cell culture, like the related Oryctes rhinoceros Nudivirus (OrNV) (37) and that these structures can be found associated with virus after viral purification. In whole animals, DiNV is primarily associated with the gut and gonads. However, gut-associated virus is mostly found in the muscle cells, not in the lumen or gut epithelium.

## Results

### DiNV capsid protein accumulates in cytoplasmic puncta but is present in both the nucleus and cytoplasm

We inoculated *D. innubila* Dinn-1 cells and stained with DAPI and immunofluorescence using an antibody (36) against the viral nucleocapsid protein (vp39) to determine gross localization patterns of the virus in cell culture. While the mock inoculated cells show no evidence of DiNV (orange staining), inoculated cells are infected (Fig 1A-B). Though nudiviruses replicate in the nucleus of cells (28), nearly all signal from the anti-vp39 antibodies was found in puncta in the cytoplasm (Figs 1B’ and 1B’’), outside of the nucleus. These puncta are reminiscent of what Velamoor *et al*. (37) referred to as multimembrane vesicles in beetle cells inoculated with Oryctes rhinoceros Nudivirus (OrNV). Finally, we anecdotally observed no consistent evidence of cytopathic effect in infected cells (Figs 1A’ and 1A’’ vs. 1B’ and 1B’’).

**Figure 1.**
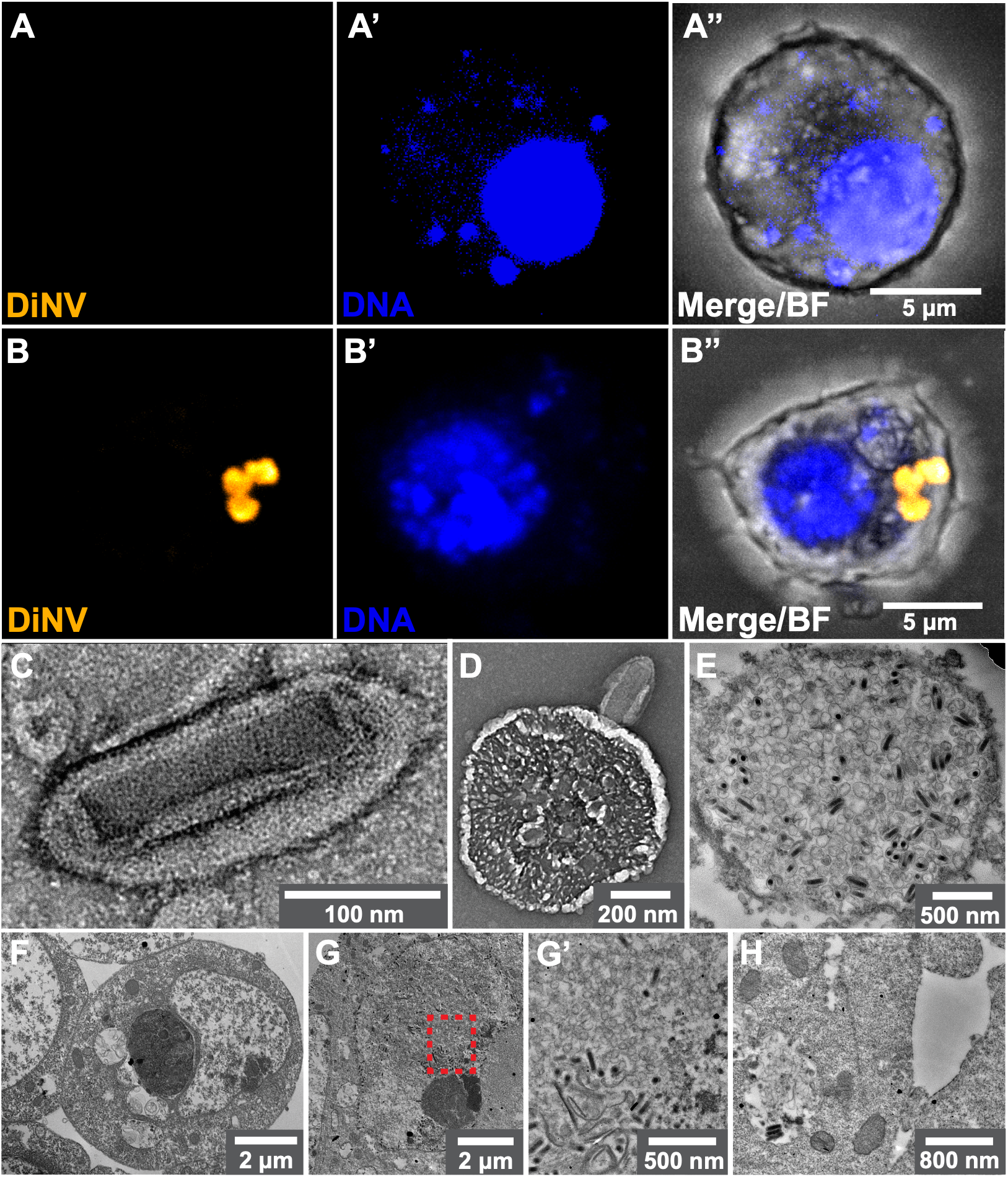
DiNV infection in cell culture and purified. A) control cell 10 days post inoculation with cell culture media with virus stained with anti-vp39 antibody (orange), nucleic acids stained with DAPI (blue), and merged with brightfield). A’ and A’’) individual cells. B) Experimentally infected cell 10 days post inoculation with DiNV (staining as in A), B’ and B’’). C) A single enveloped DiNV virion purified from cell culture imaged using transmission electron microscopy. Note the rod-shaped capsid and the ovular envelope. D) an extra cellular vesicle associated with a DiNV virion isolated from cell culture. E) A thin section of a putative vesicle containing several DiNV virions (dark rod-shaped structures). F) control cell 10 days post mock inoculation. G) DiNV-infected cell 10 days post inoculation showing several rod-shaped virions (dark structures) and G’) inset expanded view of (G) with several potentially smaller empty envelope structures. H) DiNV infected cells showing viral particles in the cytoplasm.

We next used electron microscopy to examine virus purified from inoculated *D. innubila* (Dinn-1) cell culture (Figs 1C-E) or in Dinn-1 cells (Figs 1F-H). As described previously for both DiNV and other nudiviruses (22, 28, 38), the DiNV virion contains a rod-shaped capsid (approximately 170nm by 35nm) surrounded by an envelope (Fig 1C). Curiously, several viruses in the purified sample were found associated with structures like the multimembrane vesicles described by Velamoor *et al*. (37) (Figs. 1D-E). While the virus was missing in both the mock inoculated control cell and purified samples, similar structures were present in the purified control samples (Fig S1). Figure 1E shows a section of one of the structures with several DiNV virions (dark rod-shaped capsids) but also what appear to be several empty envelopes (unfilled circles). In cell culture, the virus was absent in mock inoculated cells (Fig 1F), but clearly present in both in the nucleus (Fig 1G and 1G’) and the cytoplasm (Fig 1H) of inoculated cells. It is possible that the structure containing virions in Figure 1H is a precursor of the multimembrane vesicle and like the structures in the purified virus (Figs 1D-E), but difficult to say with certainty. We also note that much of the signal in the immunofluorescence images (Fig 1B) may either be from capsid protein not incorporated into functional viral particles or from the multimembrane vessicles. We conclude that DiNV follows an infection cycle in cells like other nudiviruses including both replication in the nucleus and packaging into multimembrane vesicles in the cytoplasm (37).

### Systemic infection of DiNV localizes to the gut and reproductive tissues

We next aimed to study viral tropism in adult flies injected with DiNV. To that end, we injected 4–7-day old males and females with DiNV and fixed in formalin 5 days after infection followed either by cryo-sectioning or dissection to examine individual tissues. Though not exclusive, DiNV primarily localized to the abdomen in both sexes, dominated by the gut and, to a lesser extent, the gonads (Figs 2A-D, Figs S2). Focusing on just the abdomen, the virus is clearly associated with the gut (arrows) and gonads (arrowhead) in females (Fig 2E, 2E’, 2E’’) and males (Figs 2F, 2F’, 2F’’). The ovary in Fig 1E appears atrophied, consistent with what was reported for Kallithea virus (32), but we did not find this atrophy consistent enough (many ovaries of infected females were unaffected) to be confident that it was associated with DiNV infection. In all tissues, DiNV appears to be associated with, but somewhat peripheral to the actual tissues. Once again, we found no evidence of virus infection in uninfected flies (Figures S2 and S3).

**Figure 2.**
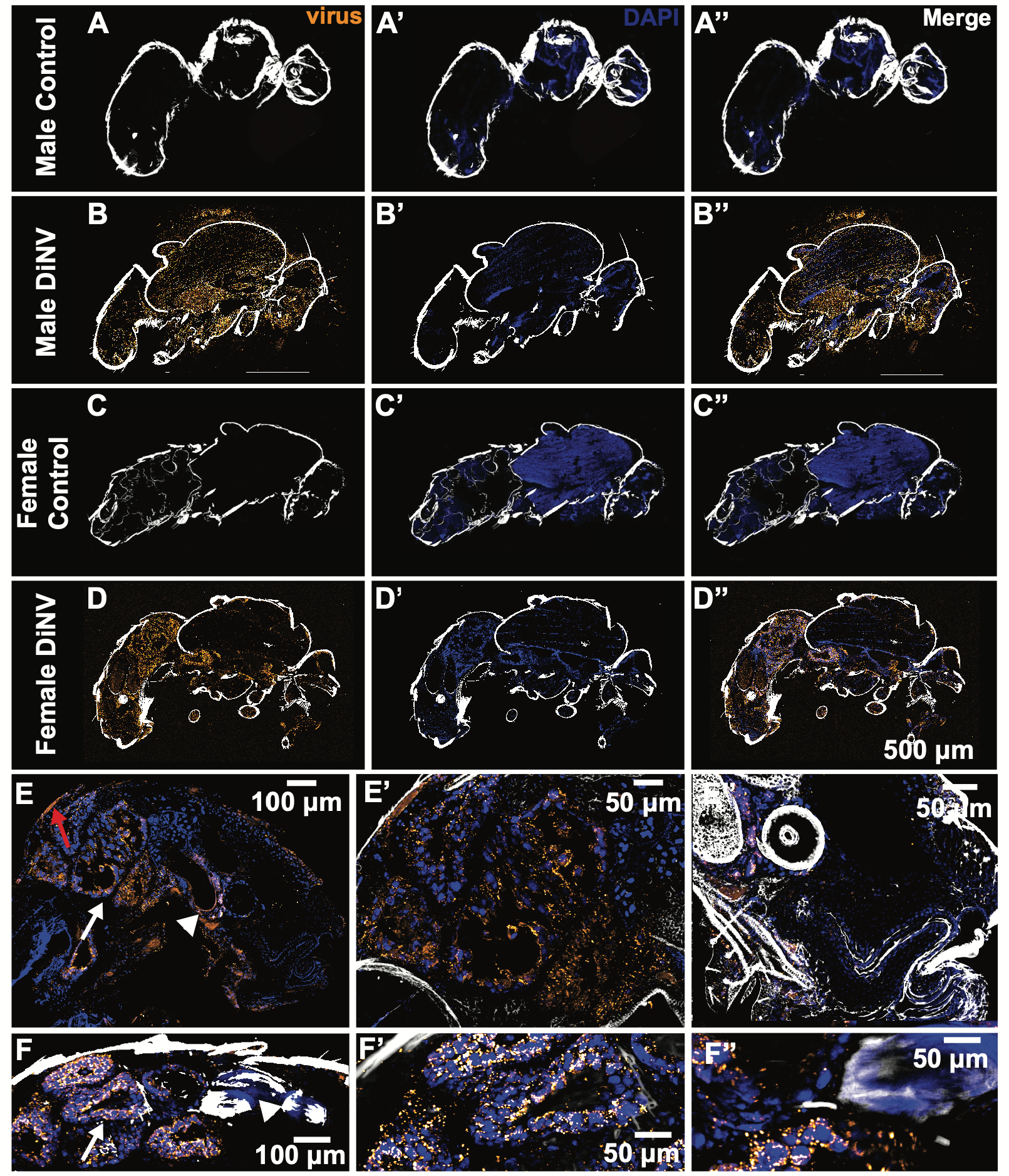
DiNV appears to localize to gut and reproductive tissue even after systemic infection. A-D) Sagittal sections of male (A, B) and female (C, D) *Drosophila innubila* either mock uninfected (A, C) or infected (B, D) with DiNV. E) Sagittal section of infected female abdomen showing gut (white arrows) and ovary (white arrowhead). F) Sagittal section of infected male abdomen showing gut (white arrows) and testes (white arrowhead). All images were stained with rabbit anti-P39 antibody and then goat anti-rabbit labeled with Alexifluor-488 but shown as orange color in the images to visualize the virus localization in the cryosections. DAPI is in the mounting medium which stains DNA and is the blue color in the images. The fluorescent signals were overlayed onto a transformed brightfield image and processed with FIJI.

### DiNV is associated with the periphery of gut tissue and mostly absent from the lumen

Many nudiviruses, and we suspect DiNV, are transmitted primarily fecal-orally. Thus, we hypothesized that since even systemic infection seemed to localize to the gut, that DiNV might leak into, and infest the lumen of the gut. We dissected midguts and hindguts from female flies, 5 days after infection and stained for nucleic acids (DAPI, blue) and DiNV nucleocapsid vp39 antibody (orange). In all samples, the midgut appeared to be highly infected with DiNV (Fig 3B), but there was almost no signal in the hindgut (top right of 3B – blue staining but no or less orange). Mock infected flies never showed any evidence of DiNV infection (Fig 3A, Figure S4). Fig 3C shows a small fragment of an uinfected midgut (approximately the third section if section one is distal to the foregut and section five is proximal to the hindgut (39, 40)). In Figure 3D, DiNV is primarily located peripheral to the stained gut nuclei – not interspersed. Figure 3F shows infected midgut cells with puncta like what were observed in cell culture (Fig 1B). Finally, a cross section of the infected gut provides further evidence that DiNV accumulates in the periphery of the gut tissue and *not* in the lumen (Fig 3H), however there is a region with more virus that may be associated with damage to the gut barrier and infiltration into the lumen. However, such instances were rare in our study.

**Figure 3.**
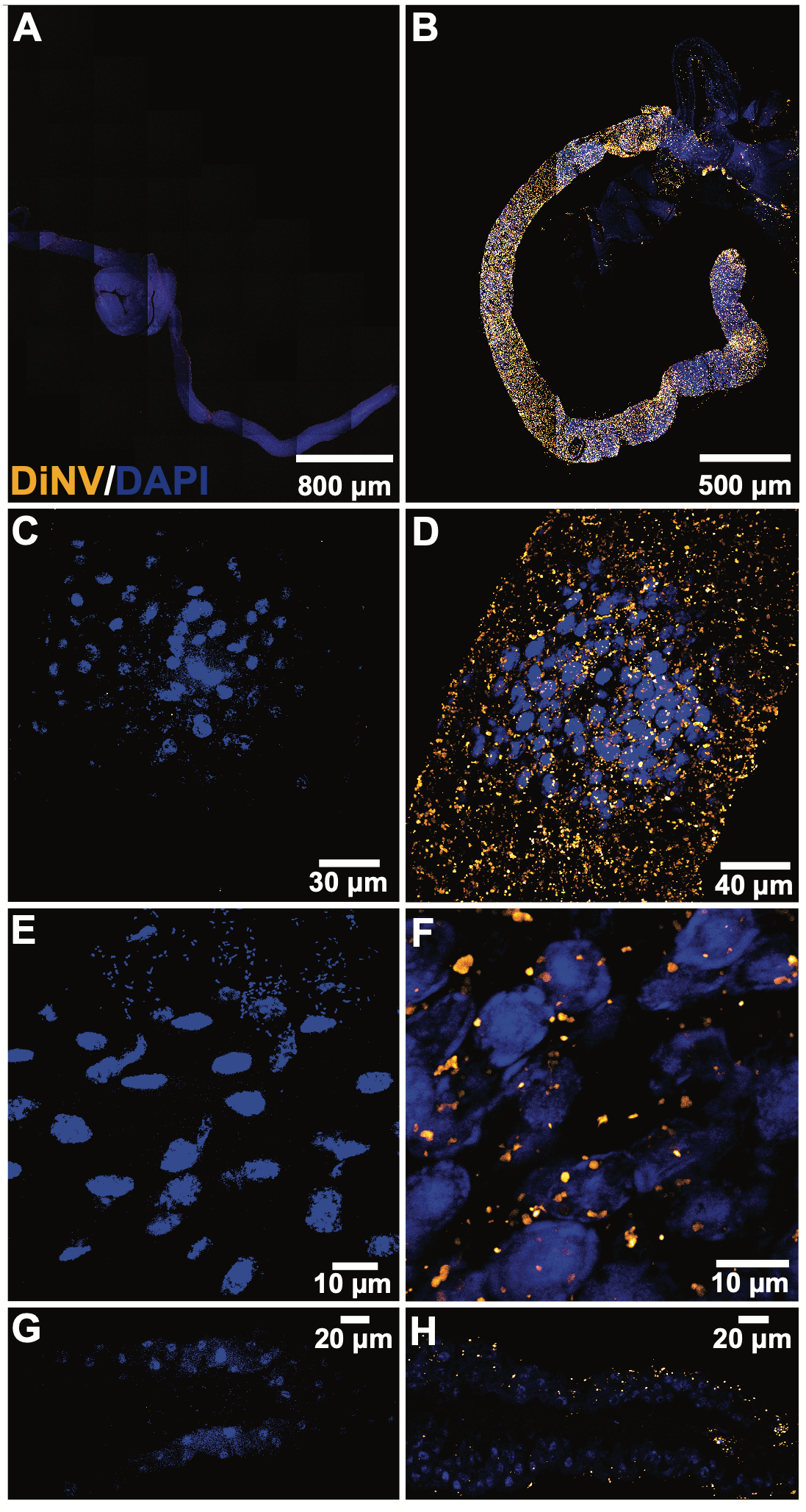
DiNV infects the gut but appears to be localized in the outer tissues. A-B) Uninfected and infected female guts dissected 5 days post infection show overwhelming signal from DiNV (orange). C) and D) are higher magnification images of A and B, and E) and F) are further magnification. G and H) Uninfected and infected female gut cross-section showing virus primarily peripheral to the gut lumen with a possible incursion of virus into the lumen.

### DiNV is primarily associated with visceral muscle cells in the gut

To further examine the localization of DiNV in the gut after systemic infection, we examined cross-sections of guts under transmission electron microscopy. Figure 4A shows the gut of an infected individual in cross section from the anterior midgut and upon close inspection, DiNV is primarily associated with the muscle cells that surround the gut, not the cells of the gut *per se*. This figure shows the peritrophic matrix orange), villi (blue), and muscle cells (maroon). Figure 4B (and 4B’) shows DiNV in the nucleus of the anterior midgut visceral muscle. DiNV infection is not exclusive to the visceral muscle and can be found in other gut-associated cells in the anterior midgut (Fig 4C). The middle midgut region (approximately R3 (39)) shows similar patterns of infection primarily in the visceral muscle (Figs 4D and 4D’). Note at least one virion outside the nucleus in the top left of Fig 4D’. We conclude that muscle cells associated with the *Drosophila* gut are especially susceptible to DiNV after systemic infection but note that DiNV *can* infect other gut-related cells.

**Figure 4.**
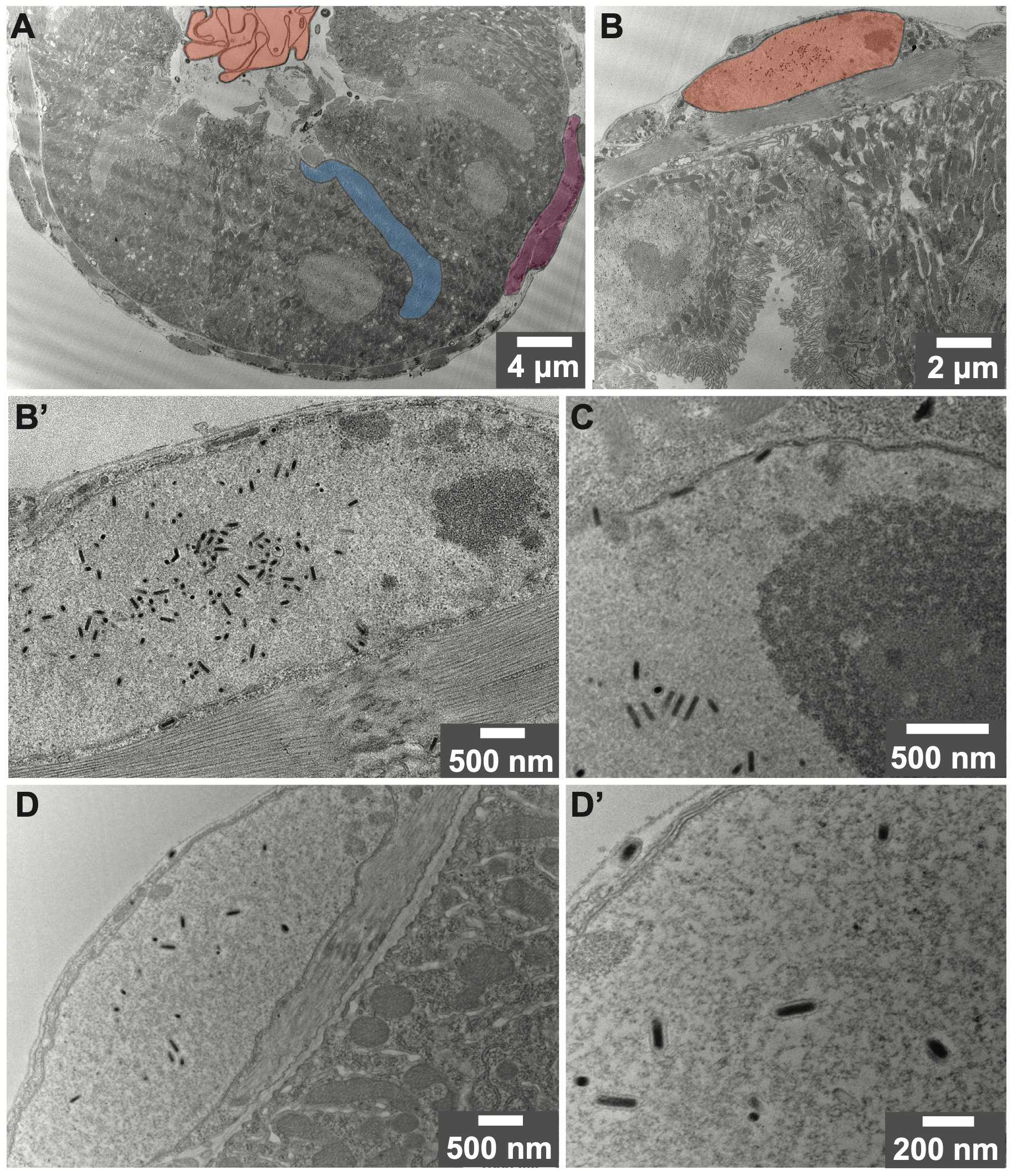
DiNV primarily infects the circular muscle cells around the midgut. A) electron microscopy of a midgut cross-section showing the peritrophic matrix (orange), villi (blue), and visceral muscle (maroon) showing virus (dark staining) associated with the visceral muscle. B and B’ show two different magnifications of a visceral muscle cell nucleus (orange in B) with significant viral presence. C) Female anterior midgut enterocyte cell showing DiNV infection in the nucleus and several virions at the nuclear membrane. D and D’) DiNV infection in the visceral muscle of middle midgut cells.

### DiNV is associated with gonads after systemic infection

As noted earlier, DiNV was associated with both ovaries and testes after systemic infection. We therefore dissected reproductive tissues to look more closely at DiNV localization in these tissues (Fig. 5, and Fig. S5). As expected, there is no evidence of infection in the mock-infected controls (Fig 5A and 5C).

**Figure 5.**
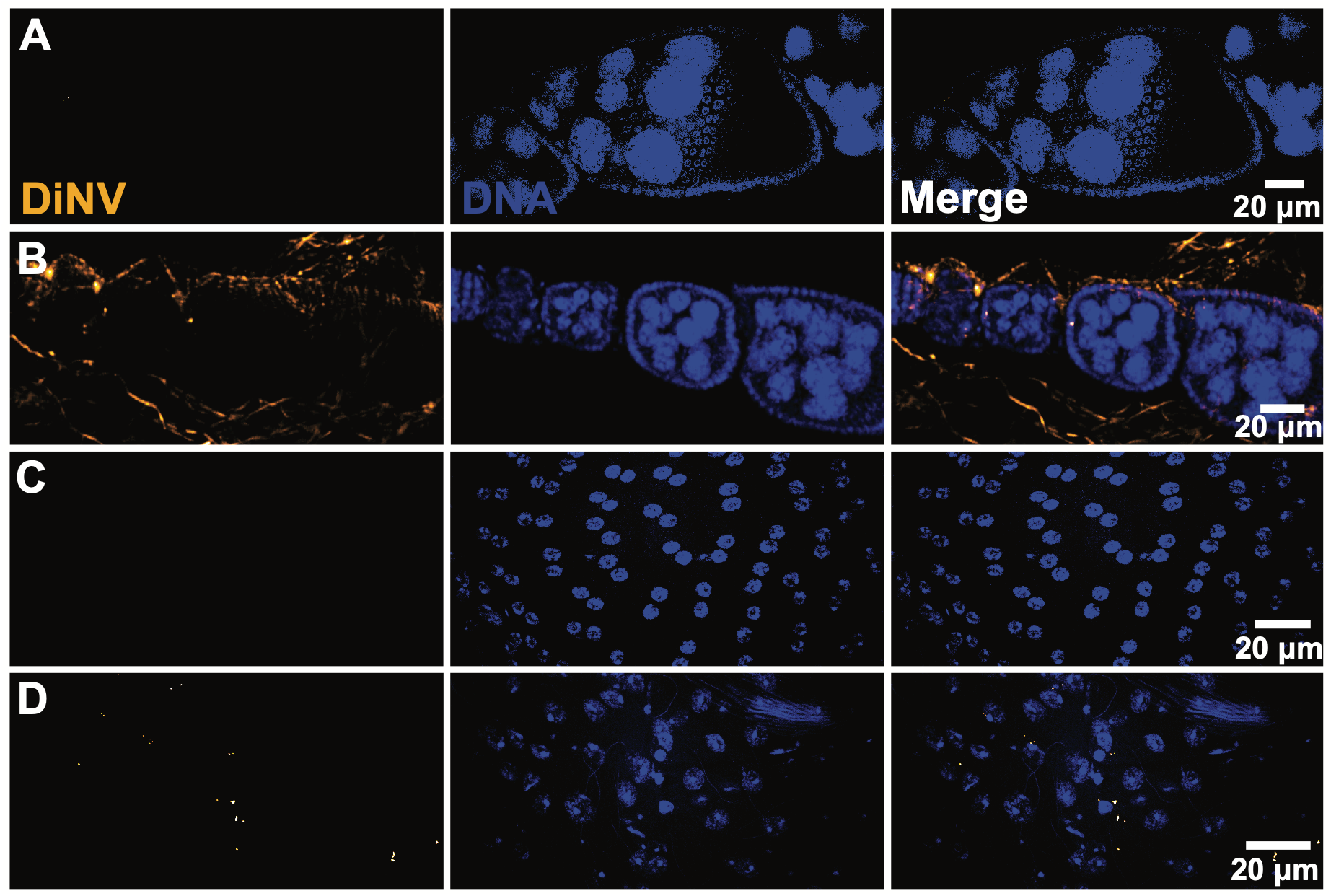
DiNV infects the reproductive tissues. Mock infected (A, C) and DiNV infected (B, D) ovarioles (A, B) and testes (C, D) 5 days post infection. Blue stain is DAPI for nucleic acids, orange is for DiNV (anti-vp39 antibody). Left column is virus alone, middle is DNA alone and right is DNA and virus merged.

Infected females have several viral-associated puncta surrounding, but not necessarily inside the ovarioles (Fig 5B). This is consistent with the ovarian atrophy associated with infection by Kallithea virus (32), but also with a *lack* of vertical transmission of DiNV from mother to offspring (22) since the virus does not seem to be inside the developing egg. DiNV in testes appears to be more integrated into the main tissue and not peripheral (Fig 5D). We do not expect paternal transmission of DiNV to offspring, but this has not been explicitly tested. This association between DiNV and the gonads could also be related to sexual transmission. We have not explicitly tested this either, but some nudiviruses including *Helicoverpa zea* Nudivirus 2 (HzNV-2) can be vertically (maternally) and sexually transmitted (41). We conclude that DiNV’s presence in the gonads may be associated with the reduced fecundity observed after both DiNV (*D. innubila* and *D. falleni*) and Kallithea virus (*D. melanogaster*) infection (22, 32).

## Discussion

We studied the cellular and tissue tropism of the Drosophila innubila Nudivirus in adult *D. innubila* and in cell culture. Consistent with other nudiviruses (37), we found that the virus appears to be associated with multimembrane vesicles that form puncta when examined under immunofluorescence. Five days after systemic infection, DiNV is most prevalent in the gut tissues but also associated with the gonads. Surprisingly, however, the gut-associated virus is not found in the lumen in the gut but primarily the visceral muscle. Thus, DiNV shows signatures consistent with both gut infections of several nudiviruses (26, 28) and association with reproductive tissues like HzNV-2 (30).

Nudiviruses are sister to the baculoviruses and share a common morphology and about 20 core genes (28, 38). The formation of occlusion bodies, a protein matrix that contains one to several virions and protects the virus from degradation in the environment between shedding or death of one host and acquisition by the next host, is a hallmark of baculovirus infection (42, 43). Two characteristic structures associated with occlusion bodies are the polyhedrin protein matrix and the PIF complex, a series of 8-10 proteins that together are essential for cell entry in oral infection of occluded baculoviruses (44). Nudiviruses were named for their lack of this occlusion body, but the reality is more complex. Most nudivirus genomes contain genes encoding both polyhedrin and multiple PIF complex components (38, 44). Furthermore, several nudiviruses can form occlusion bodies, including Oryctes rhinoceros Nudivirus, a close relative of DiNV (38, 45-47). In contrast, DiNV (as in a prior study with OrNV (37)) is sometimes associated with multimembrane vesicles both in and outside of cells (Fig 1). These structures share very little resemblance to the concrete-like nudivirus or baculovirus occlusion bodies other than the accumulation of virions within. So, it is possible that the multimembrane vesicles observed in OrNV and DiNV perform a similar function to the occlusion bodies, but this fails to explain the maintenance of polyhedrin and PIF complex genes in these nudiviruses. Another possibility is that occlusion bodies are formed much later during nudivirus infection, and that our studies failed to find them because our time-courses were too short. In another study, we found that an antibody for PIF-3 a PIF complex component, is not detectable *via* Western blot in cell culture until much later (14 days post inoculation) and that the PIF complex genes are mostly unexpressed during the first five days after infection of animals or inoculation of cells (Adewumi, *et al*., in prep). On the other hand, virus detected in *D. innubila* feces was not in occlusion bodies suggesting that DiNV *can* exist outside the host as naked (but enveloped) virions (22). Finally, systemic injection of even low doses of DiNV kills *D. innubila* in about 10 days, so this infection route likely will not allow production of occlusion bodies if they take more than 10 days to develop. It remains to be seen whether an oral infection leads to slower rates of death and therefore the possibility of occlusion body formation.

DiNV quickly migrates to the gut and gonads after systemic infection. This suggests a relatively strong tissue tropism, but two contrasting but not mutually exclusive hypotheses about DiNV transmission dynamics. Most nudiviruses and baculoviruses are assumed to be transmitted either fecal-orally or *via* ingestion of virus associated with dead hosts (28, 38, 48). The overwhelming tropism toward the gut is consistent with fecal-oral transmission of DiNV. However, in our limited sample, the virus was mostly absent from the lumen of the gut and common in the surrounding visceral muscle. This would presumably limit the ability of DiNV to shed and be transmitted through the feces. The abundance in the visceral muscle may be a consequence of the route of infection – systemic injection likely leads to infection from the periphery of the gut inward whereas oral infection would be the opposite. So, it is possible that if we had harvested fly guts at a later timepoint, we would have seen more DiNV in the lumen, and therefore more shedding. We did find, anecdotally, that the guts of infected flies were much more fragile than those of uninfected flies and that they seemed to get progressively worse with the number of days post infection. Overall, the tropism results are consistent with transmission *via* oral infection, but additional studies are necessary to show this conclusively.

DiNV was also associated with both testis and ovary tissue (Fig 5), though to a qualitatively lesser extent than in the gut. Interestingly in both testis and ovary, DiNV was peripheral to the gametes, not directly incorporated (Figs 2E, 2F and 5). We anecdotally noted that ovaries of infected females were smaller and contained fewer eggs as was found for Kallithea virus (32), but hesitate to speculate on whether this is a consequence of DiNV infection, and if so whether it is due to a general reallocation of resources away from reproduction and toward immune defense or a direct consequence of DiNV’s association with the ovary. HzNV-2 is sexually transmitted and causes infected individuals to be effectively sterile (30, 41, 49), but also quite distantly related to DiNV among the nudiviruses (33). Overall, there is incomplete evidence for both oral acquisition of DiNV through the gut and sexual transmission. Both routes of transmission are possible, and the availability of multiple transmission routes would likely enhance the fitness of DiNV.

## Materials and Methods

### Viral culture, cell lines and fly stocks

All the virus used in this experiment for inoculation or infection was isolated from *Drosophila innubila* (Dinn-1) cells inoculated with cell-culture adapted virus, incubated for 10 days, frozen, then thawed and filtered through a 0.45-micron filter (36). In contrast, the purified viral material examined by electron microscopy was from the supernatant of the clarified pellet of a purification protocol (36). Dinn-1 cells were used for all cell-culture experiments and were generated in our lab by homogenizing embryos and culturing until established and capable of freeze/thaw cycles and passaging (36). The *D. innubila* genome strain was used for infection (50). They are kept on a 12hr light/12hr dark cycle at 20C without humidity control. Their media is a teaspoon of instant Drosophila media (Carolina, USA) hydrated and supplemented with a chunk of store-bought mushroom (*Agaricus bisporus*) and a cotton roll (51).

### Immunofluorescent Confocal Microscopy

All confocal images were obtained using an Olympus FV1000 laser-scanning confocal microscope (Olympus America Inc., Center Valley, PA, USA) equipped with Fluoview software. Z-stacked images were collected and exported to FIJI (52). We collected images using the 488 nm filter, and DNA using the DAPI filter. Along with the 488 nm, DAPI and a brightfield signal we collected a signal from the 546 nm laser to subtract autofluorescence from the 488 and DAPI signals using the FIJI Image Calculator. For images the 488 nm signal was set to orange and DAPI to blue. The brightness and contrast of each image was adjusted and then merged to create the overlay images.

### Imaging DiNV infected cells with immunofluorescence

Dinn-1 cells were seeded onto two Geltrex (A1413202, GIBCO, Jenks, OK, USA) coated Lab-Tek Flaskette Chamber Slide system (Flask style with Glass slide, 177453, Thermo Scientific, Waltham, MA, USA) in Schneiders Insect Medium with heat inactivated FBS (10% final concentration) and gentamicin (10μg/ml). The cells were incubated at 23°C for 14 days, then fluid changed to Schnieder’s medium with gentamicin and then 50 μL of adapted virus (36) was added to the culture medium. A second flask was mock inoculated with growth medium. Cells were incubated at 23°C for 10 days when the media was removed, and the cells were fixed with 2 mL of 2% paraformaldehyde (PFA) for 30 minutes. After fixation the PFA was removed, and the slides were rinsed 3 times with DPBS. To minimize nonspecific antibody binding we treated the cells with DPBS with BSA (2% final concentration) for 30 minutes. Next, the slides were incubated with a rabbit polyclonal antibody with specificity to the DiNV nucleocapsid (anti-P39) diluted in DPBS for 1 hour at room temperature. The cells were washed again and incubated with Alexa fluor 488 goat anti-rabbit IgG (Invitrogen) for 1 hour at room temperature followed by a final wash (3x) with DPBS. Finally, the cells were overlayed with Vectasheild Vibrance Antifade Mounting Medium with DAPI (Vector Laboratories, Newark, CA, USA) and covered with a coverslip and stored flat at room temperature in the dark.

### Imaging sagittal cryosections of infected flies for confocal microscopy

Adult male and female flies were injected with DiNV as described previously (36). Briefly, 5-7 day old adults were injected by nanoinjector into the thorax with 27.6 nL of viral fluids, then transferred to new food and observed. After 5 days the infected and control flies were anesthetized with CO_2_ and fixed in 2% paraformaldehyde in DPBS (21-031-CV, Corning Life Sciences, Tewksbury, MA, USA) for 1 hour. The flies were then transferred to 12% sucrose in DPBS and held at 4°C for 24 hours, then transferred to 20% sucrose in DPBS followed by incubation at 4°C. After 24 hours the flies were embedded in Tissue Tek (OCT) using a cryo-tissue holder. The embedded flies on chucks were stored at -20°C until sectioning. The cryostat temperature was set to -22°C and the section thickness to 20 μM. Cut sections were carefully placed on gelatin subbed slides (slides were equilibrated to -22°C). The slide with the section was brought to room temperature and gently heated with a touch of fingertip to the underside of the slide to melt the OTC matrix. The slides were then dried by incubating at 37°C for 30 minutes. The dried sections on slides were stored at -80°C until staining.

Slides were thawed and submerged in 2% PFA for 15 minutes. The OTC dissolved away from the tissue and the PFA treatment linked the tissue section to the gelatin subbed slide. To inhibit autofluorescence of the sections the slides are treated with cold sodium borohydride (0.1% in DPBS) for 5 minutes with 3 changes. After the sodium borohydride treatment, the slides were washed 3 times in DPBS. For the blocking step, sections were incubated in buffer consisting of 1% BSA, 20mM glycine, .05% Tween-20 in DPBS for 1 hour. After blocking the tissue was soaked in rabbit anti-vp39 diluted in DPBS containing 1% BSA and 0.1% Tween-20 for 1 hour at room temperature followed by 3 washes in DPBS. The slides were then incubated in goat anti-rabbit, Alexa fluor 488 diluted in DPBS, 1% BSA, 0.1% Tween-20 and incubated for 1 hour at room temperature then washed 3 times with DPBS. Next the slides were removed from the last wash and the liquid around the slide was blotted away without touching the tissue. Lastly, 100ul Vectashield with DAPI was pipetted onto the slide and a cover glass was overlayed onto the sections.

### Imaging whole mount or dissected animals with immunofluorescence

Flies were dissected under a dissecting microscope in PBS and immediately transferred to a well of a 96 well plate containing 2% paraformaldehyde in PBS and sucrose and fixed for 1 hour at room temperature. The fixed tissue was transferred to a 70 μM cell strainer (22363548, Fisher Scientific, Waltham, MA, USA) that was placed in a well of a 6 well plate containing DPBS. The tissue was washed 3 times with DPBS to rinse out the fixative followed by a 15-minute soak in DPBS with Triton (0.1%) then washed 3x in DPBS. Staining for DiNV and DNA followed the same procedure described above. The slides were kept in the dark at room temperature for 24 hours to allow time for the mounting medium to solidify then viewed by fluorescent confocal microscopy.

### Imaging cells and purified virus samples with electron microscopy

Dinn-1 cells growing in 25 cm^2^ plug seal flasks were inoculated with 100 μL of DiNV and incubated for 10 days. The growth medium was then removed and 10 mL of 4% formaldehyde diluted in Schneiders Medium was added back. The cells were gently scraped from the flask and the fluids were pipetted to a 15 mL conical centrifuge tube. The fixed cells were stored at 4°C. The prepared virus samples were submitted to The University of Kansas, Microscopy and Analytical Imaging Lab for preparation and observation by Electron Microscopy.

Cells in 15 ml centrifuge tubes were centrifuged and 300 μL of the pellet was moved to a new vial and rinsed twice in HBSS/0.1% sucrose (HBSS/Su) for 5 minutes. The sample was then permeabilized twice in HBSS/Su + 0.1% saponin solution (HBSS/Su/Sap) for 5 minutes, then blocked in HBSS/Su/Sap + 0.1% Triton X-100 solution + 3% Normal Goat Serum for 30 minutes at room temperature. Samples were further prepared for immunogold staining, but that did not work so is not further described here. The samples were post-fixed with a modified Karnovsky’s fixative 1 (2% formaldehyde + 2.5% glutaraldehyde) diluted in HBSS/Sucrose and incubated overnight at 4 °C. The samples were then rinsed twice in 0.2M Cacodylate buffer with sucrose for 5 minutes, stained with 2% Osmium tetroxide diluted in 0.2M Cacodylate buffer with sucrose for 2 hours at room temperature. The samples were then rinsed in 50% EtOH three times for 5 minutes each, then rinsed in 70% EtOH 5 minutes. The samples were treated with 4% uranyl acetate in 70% EtOH at room temperature for 15 minutes. Finally, they were embedded in Durcupan ACM Epoxy (mix 2, Electron Microscopy Sciences, Hatfield, PA, USA) for 48 hours at 65°C. Sections and Fluids were added directly to grids for transmission electron microscopy of 70 nm sections, and virus fluids.

Embedded cells were sectioned at 50-70 nm using a Leica UC6 ultramicrotome (Leica Microsystems, Wetzlar, Germany) with an ultra 45° angle, 2.1 mm diamond knife (DiATOME USA, No. MF 2407, Hatfield, PA, USA). Sections were collected on 100 mesh copper grids and imaged on a Hitachi H-8100 Transmission Electron Microscope (Hitachi High-Tech, Schaumburg, Il, USA) with a lanthanum hexaboride (LaB6) electron gun and an AMT BioSpirit 16 mega-pixels CCD camera (4896 x 3264 pixels, Woburn, MA, USA). All electron microscopy workflow development, sample preparation, processing and imaging were performed in the Microscopy and Analytical Imaging Resource Core Laboratory at The University of Kansas.

## Acknowledgements

We thank members of the Unckless lab especially Ananda Kalukin and Taiye Adewumi for their helpful comments and suggestions. We also thank the KU Microscopy and Analytical Imaging Research Resource Core for working with us to acquire images. Funding for this work was provided by NSF grant 2135167 to RLU and NIH grant R03-AI193975 to RLU.

